# Salivary Biomarkers for Classifying Acute Physical Fatigue: A Comparative, Exploratory Study

**DOI:** 10.1101/2025.06.04.657971

**Authors:** Bryndan Lindsey, Kenneth Bowden, Yosef Shaul, Emanuel Petricoin, Shane V. Caswell, Rayan Alhammad, Anissa N. Elayadi, Brandon M. Roberts, Joel R. Martin

## Abstract

**Introduction:** Physical fatigue is a key determinant of operational readiness in tactical athletes. Hormonal, immune, and enzymatic biomarkers have been proposed for fatigue assessment, but reliability can be affected by external factors. Therefore, this study aimed to compare targeted stress-related biomarkers, proteins identified via untargeted salivary proteomics, and a combined panel of both for classifying acute physical fatigue.

**Methods:** Ten recreationally active adults (6M, 4F) completed a fatiguing protocol within a 3-day sample collection window. Saliva samples were analyzed for targeted biomarkers via commercial immunoassays and proteins via untargeted liquid chromatography-mass spectrometry. Targeted, protein, and combined panels were identified using nested leave-one-subject-out cross-validation with FDR correction and stability selection, then classified via ridge-penalized logistic regression. Models were extended across the full protocol to evaluate classification of fatigue onset, rest, and recovery states.

**Results:** A candidate identified protein (C1S) achieved slightly higher classification accuracy (80%) relative to a panel of targeted biomarkers (IgA and UA 70%), with superior sensitivity but similar specificity. In comparison, the combined panel, spanning immune (IgA), oxidative/metabolic (UA), complement (C1S), and cellular stress-related (ANG) outperformed both single-platform models with 90% accuracy, sensitivity, and specificity.

**Conclusions:** Improved classification accuracy observed with the inclusion of candidate salivary proteins provide preliminary support for non-invasive proteomic monitoring of fatigue in tactical settings, and in particular the combination of complementary targeted and proteomic panels to capture a more complete physiological signature of acute fatigue. Future studies should validate these findings in larger, more diverse populations and assess applicability for chronic fatigue monitoring.

## Introduction

Operational readiness, commonly described as the ability of military forces to fight and meet the demands of the assigned missions, is shaped by various factors (1, 2). One important factor is fatigue, which is characterized by a measurable decline in muscular performance or an increase in perceived exertion (3) that can impair functional capacity, elevate injury risk, and compromise mission success (4). Following sustained occupational activities, fatigue is frequently elevated; however, capturing this in a rapid, objective manner remains challenging, especially in tactical populations.

Physical exertion induces coordinated physiological stress responses that are reflected in measurable changes in circulating biomarkers (5). Specifically, prolonged physical exertion causes changes in the autonomic nervous system (ANS), influencing the activation of hormones like cortisol and testosterone amongst other compounds (6). Biomarkers such as cortisol and testosterone (7), immunoglobulin A (IgA) (8), alpha-amylase (AA) (9), interleukin-6 (IL-6) (10), and uric acid (UA) (11) have all been shown to change as a result of military related stressors, and therefore, have been proposed as potential biomarkers of physical readiness. However, to date such biomarkers have predominantly been quantified via blood samples which is invasive, requires trained personnel, and presents compliance and logistical challenges for repeated sampling in operational settings.

Saliva sampling offers a novel, non-invasive, easily collected medium for assessing biomarkers rapidly and in field environments, with recent evidence suggesting that salivary biomarkers are responsive to a variety of acute stressors like muscle damage, sleep deprivation, and dehydration (12). Recently, untargeted proteomic approaches have identified salivary proteins related to immunoregulation, inflammation, and metabolism that are associated with fatigue status in civilian (13, 14) and military settings (15). These proteins have been hypothesized to be more specific markers of physical fatigue as opposed to hormones like cortisol and testosterone which are influenced by diurnal rhythm and psychological stressors (16). However, the comparative discriminatory performance of proteomic biomarkers relative to established stress-related salivary biomarkers has not been directly compared within the same cohort. Additionally, combining complementary assay platforms may improve the characterization of fatigue beyond either approach alone, consistent with broader evidence that integrating complementary biomarkers across molecular pathways improves signature detection (4, 15).

Therefore, this exploratory study aimed to identify candidate biomarkers of acute physical fatigue from targeted stress-related biomarkers and salivary proteins identified via untargeted liquid chromatography-mass spectrometry (LC-MS), and to compare the relative classification accuracy of single-platform and combined biomarker panels for detecting acute physical fatigue, following a controlled fatiguing protocol.

## Methods

### Study Participants

Ten recreationally active adults (6 males, 4 females; age 32.4 *±*9.1 years) were recruited from the kinesiology department of a public university in the United States between April 1, 2023 and August 30, 2023. This convenience sample comprised physically active individuals with a range of fitness levels, training backgrounds, and exercise experience. Inclusion criteria included ages 18–55, absence of injury or chronic disease, and ability to complete a weighted 2+ mile ruck. Participants engaged in ≥30 minutes of daily physical activity and reported no limitations in performing standard exercises. All provided written informed consent that was witnessed by the study principal investigator. The study was approved by the university’s IRB (IRB #2004820) and conducted in accordance with the ethical standards set by the Helsinki Declaration.

### Data Collection Protocol

Saliva samples (*∼* 5mL per collection) were collected at 7 timepoints over a 3-day span (Figure 1) using the mLife True oral fluid collection device (mLife, TX, USA). The protocol spanned three days: baseline (day 1), fatigue (day 2), and recovery (day 3). On the baseline and recovery days, samples were taken at approximately 0800 (Measure 1) and 1300 (Measure 2) hours, and participants were instructed not to perform any strenuous exercise 24 hours before day 1 or during the protocol outside the prescribed fatiguing protocol on day 2. On the second day, participants performed an hour-long physical fatiguing protocol, donning a wearable electro-cardiogram (ECG) monitor (Zephyr Bioharness 3.0, Zephyr Technology Corporation, Annapolis, MD, USA) which they wore approximately 1-hour before, during, and 1-hour after the protocol.

**Fig. 1.**
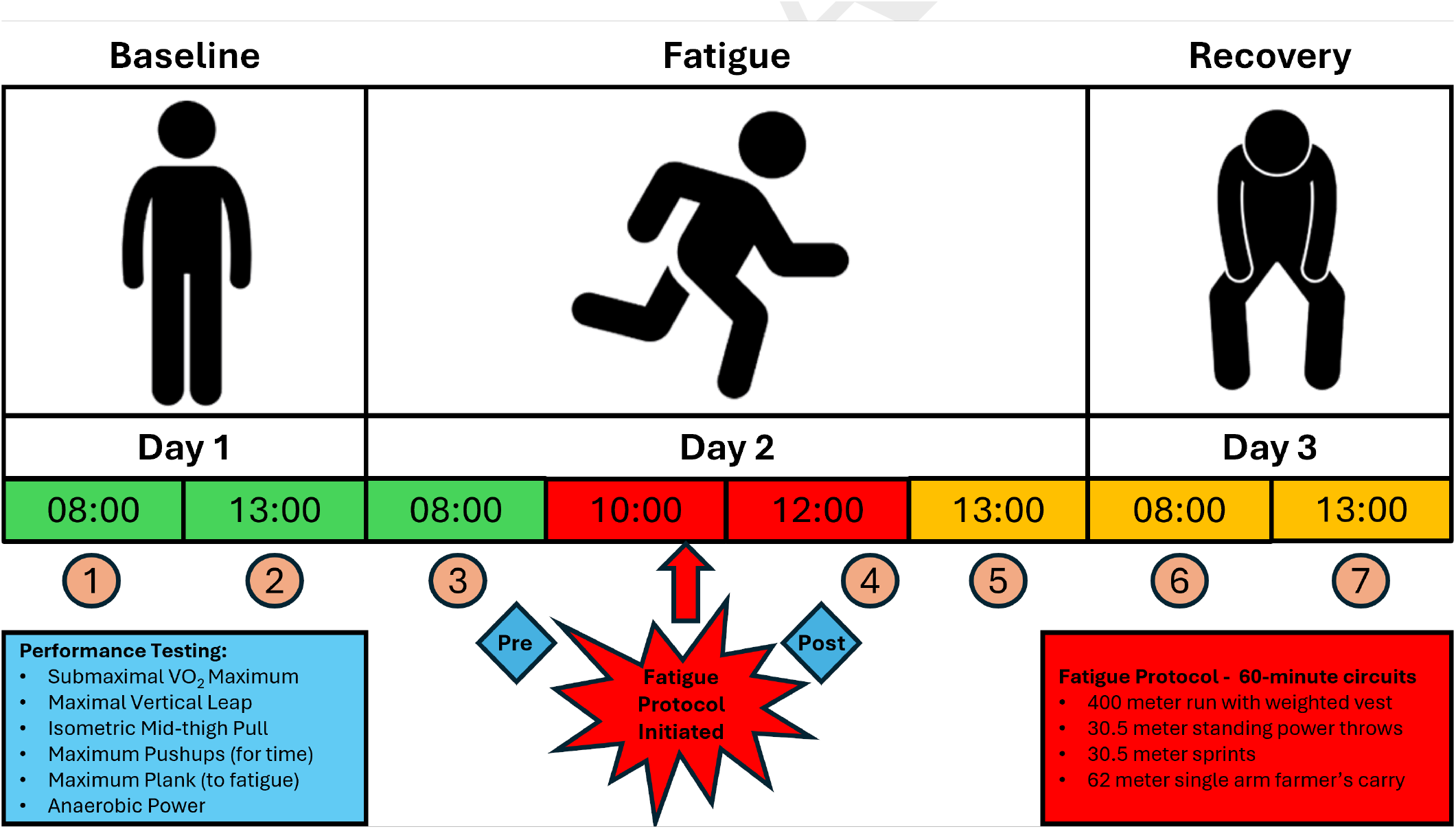
Overview of the three-day experimental protocol and saliva sampling timeline. Baseline measurements were collected on day 1, followed by a fatigue induction protocol on day 2 and recovery monitoring on day 3. Saliva samples were obtained at standardized time points (approximately 08:00 and 13:00) during baseline and recovery, as well as before (Measure 3), immediately after (Measure 4), and one hour following (Measure 5) the fatigue protocol on day 2. The fatigue protocol consisted of a 60-minute circuit of running, sprinting, power throws, and loaded carries, followed by a battery of physical performance tests assessing aerobic, neuromuscular, and anaerobic function. Numbered circles denote the order of saliva sample collection.

Pre- and post-fatigue heart rate (HR) and heart rate variability (HRV), the latter computed as a rolling 300-beat standard deviation of normal-to-normal R-R intervals (SDNN) values (17), were processed using the Zephyr Bioharness 3.0 proprietary algorithm and calculated as 30-minute averages immediately before and after completion of the protocol. Both durations represented periods of quiet sitting where participants provided saliva samples while answering pre- and post-fatigue surveys. The full period of quiet sitting was used to quantify HRV, as signal stability has been shown to increase with longer recording durations (18) while the quiet, controlled conditions minimized the likelihood of motion artifact, noise, and ectopic beats that could otherwise compromise signal quality.

The fatigue protocol involved participants performing one hour of continuous circuits, each including a 400m run, 30.5m standing power throw (4.5kg), 30.5m sprint, and a 62m single arm farmer’s carry (M = 22.7kg, F = 15.9kg) while wearing a weighted vest (9.1kg). This circuit was developed to simulate common occupational tasks and movements in operational settings. Directly before and after the fatiguing protocol, participants performed a series of physical performance tests which included Queen’s step test (19), maximum standing vertical leap (CMJ) and isometric mid-thigh pull (IMTP) on force plates (ForceDecks, Vald Performance, Brisbane, Queensland, Australia), maximum hand-release pushups (HRPU) and plank hold, and average anaerobic power (watts) over a 60 second maximal sprint on cycle ergometer (max power; Rogue Echo Bike, Rogue, Columbus, Ohio, USA). On the fatigue day (day 2), saliva samples were collected at approximately 0800 (Measure 3), 1200 (Measure 4; directly after completion of the post-fatigue protocol performance assessment) and 1300 (Measure 5; one hour after the post-fatigue protocol performance assessment). On the recovery day (day 3), participants returned to the lab to have saliva samples collected at approximately 0800 (Measure 6) and 1300 (Measure 7). Samples were stored on dry-ice until cessation of each data collection day after which they were preserved at −80^*°*^C at the Center for Proteomics and Molecular Medicine (CAPMM) until analysis.

### Targeted Biomarker Analysis

Aliquots of saliva (*∼*0.5 mL) were analyzed for cortisol, testosterone, IgA, AA, IL-6, and UA via immunoassays at a commercial lab (Salimetrics, Carlsbad, CA). Before testing, samples were thawed to room temperature, vortexed, and centrifuged at approximately 3,000 RPM (1,500 *×*g) for 15 minutes. High-sensitivity enzyme immunoassays were used to quantify cortisol, testosterone, and IgA, while kinetic enzyme immunoassays were employed for AA and UA. IL-6 was measured using the Salimetrics Cytokine Panel and assayed in duplicate at the Salimetrics SalivaLab (Salimetrics LLC, State College, PA, USA) using a proprietary electrochemiluminescence method developed and validated for saliva. Mean values of duplicate measures of targeted biomarkers were used for all subsequent analyses.

### Proteomic Analysis

The remaining 4.5 mL from each saliva sample was analyzed at the Center for Applied Proteomics and Molecular Medicine (CAPMM) at George Mason University to identify and quantify salivary proteins. Low abundance proteins were enriched by selective capture using core-shell hydrogel nanoparticle based technology as previously described (20, 21). After multiple washes to remove unbound proteins, the protein-nanoparticle complexes were eluted by incubation with 4% sodium dodecyl sulfate (SDS) in 50 mM ammonium bicarbonate at room temperature. The SDS was then removed using detergent removal columns according to the manufacturer’s instructions.

Eluted proteins were digested with trypsin at 37^*°*^C for 6 hours overnight, followed by quenching with the addition of 2 *µ*L of Trifluoroacetic Acid (TFA). Trypsinized peptides were purified using C18 spin columns with 80% acetonitrile/0.1% formic acid solution. Liquid chromatography-tandem mass spectrometry (LC-MS/MS) was performed using an Exploris 480 mass spectrometer (ThermoFisher Scientific, Waltham, MA, USA) equipped with a nanospray EASY-nLC 1200 HPLC system. Peptides, resuspended in 0.1% formic acid were separated using a reversed-phase PepMap RSLC C18 LC column (ThermoFisher Scientific) using a mobile phase of 0.1% aqueous formic acid (A) and 0.1% formic acid in 80% acetonitrile (B).

Data acquisition was performed in data-dependent mode, with one full MS scan (300–1500 m/z, 60,000 resolving power) followed by MS/MS scans of the most abundant molecular ions, dynamically selected and fragmented by higher-energy collisional dissociation (HCD) at a collision energy of 27%. Tandem mass spectra were searched against the NCBI human database using Proteome Discover v 2.4 (Thermo Fisher Scientific). Database searches employed the SEQUEST node with full tryptic cleavage constraints and dynamic methionine oxidation. Mass tolerances were set to 2 ppm for precursor ions and 0.02 Da for fragment ions. A 1% false discovery rate (FDR) was applied to peptide-spectrum matches (PSM) as the reporting threshold. Precursor Ions quantifier node was used to for protein abundance calculation and quantification.

### Statistical Analysis

#### Performance Parameters

As all participants completed the same fatiguing protocol rather than one scaled for individual fitness, changes in performance tests and cardiac metrics were used to verify the induction of physical fatigue. Normality was assessed using the Shapiro-Wilk test, and paired Student’s t-tests were used to compare pre- and post-fatigue measures (*p <* 0.05), with effect sizes calculated using Hedges’ *g* for paired samples.

#### Identification of Candidate Targeted Biomarkers

Transformed concentrations of all six targeted biomarkers (cortisol, testosterone, IgA, AA, UA, and IL-6) were filtered to Measure 3 and 4, representing the pre- and immediately post-fatigue timepoints, respectively. Final targeted biomarker panel selection and classification were performed with a ridge-penalized logistic regression with a nested leave-one-subject-out cross-validation (LOSOCV) framework. At each of 10 folds, one participant’s data were held out, and mixed-effects models were refit on the remaining 9 participants to identify biomarkers showing only a significant fatigue response (Measure 3 vs. 4) independent of diurnal variation (Measure 1 vs 2); fatigue-response p-values were corrected for multiple comparisons within each fold using the Benjamini-Hochberg (BH) false discovery rate (FDR) procedure (*q <* 0.05). Z-score standardization parameters were likewise derived exclusively from the training participants and applied to the held-out participant. Because the retained biomarker set could vary across folds, we reported the proportion of folds in which each biomarker was retained with the final targeted biomarker panel consisting only of those retained in at least 80% of folds (22). This threshold, combined with within-fold FDR correction, was designed to balance sensitivity to genuinely fatigue-responsive biomarkers against the false-discovery risk of testing multiple candidates in a small, exploratory sample.

#### Identification of Candidate Proteins

Untargeted mass spectrometry of our samples yielded 2,284 proteins across time points. Of the 2,284 proteins detected, those present in more than 50% of samples (*n* = 1,352) were retained. Across the 1,352 candidate proteins and 69 samples (93,288 total protein-sample values), 11,450 values (12.3%) required imputation; 851 proteins (63.0%) had at least one missing value, 541 (40.0%) exceeded 10% missingness, and 374 (27.7%) exceeded 20% missingness, with a maximum missingness of 49.3% for any single protein. Missing values were imputed at 50% of the detection limit, and all data were log-normalized.

Similar to the selection and classification of fatigue using targeted biomarkers, protein screening, panel selection, and classification (ridge-penalized logistic regression) were performed using the same nested LOSOCV framework. At each of 10 folds, one participant was held out, and the remaining 9 participants’ data were used to: (1) narrow the 1,352 prevalence-filtered proteins to the most reliable candidates by rested-phase (Measures 1–3) measurement consistency, retaining the most consistent 50% to reduce the number of proteins entering significance testing and the associated false-discovery risk; (2) fit linear mixed-effects models for each retained protein, with timepoint as a categorical fixed effect and participant as a random intercept, retaining proteins demonstrating a significant fatigue response (Measures 3 vs. 4, BH FDR corrected within each fold, *q <* 0.05) independent of diurnal variation (Measure 1 vs. 2); and (3) extract the acute fatigue mixed-effects coefficient for each retained protein as an effect-size estimate, selecting proteins with an absolute effect size *≥* 2.0 log units as the fold-specific protein panel. Z-score standardization parameters were derived exclusively from the training participants and applied to the held-out participant. Similar to targeted biomarkers, the proportion of folds in which each protein was retained was reported, and proteins were required to be retained in 80% of folds to be included in the final protein panel.

#### Imputation Sensitivity0020Analysis

To assess the robustness of protein selection to the choice of missing-data imputation method, the full-sample mixed-effects screening and BH FDR-corrected significance criterion described above was repeated under two additional detection-limit substitution fractions and compared against the primary approach (50% of the protein-specific detection limit): 75% of the detection limit and 100% of the detection limit, each testing sensitivity to the specific substitution fraction chosen under the same left-censored, missing-not-at-random assumption. For each scheme, the number of proteins meeting the BH-corrected fatigue-significance screening criterion, their overlap with the primary screening result (Jaccard index), and the identity of the resulting protein list were compared.

#### Identification of Combined Candidate Biomarker Panel

To evaluate whether combining targeted and proteomic biomarkers improved classification of acute fatigue beyond either platform alone, targeted biomarkers and proteins were screened jointly within the same nested LOSOCV framework, rather than combining each modality’s separately-identified panel post hoc. At each fold, all 6 targeted biomarkers and 1,352 proteins (1,358 candidates total) were screened together on the 9 training participants using the same fatigue-response criterion described above (BH FDR corrected within each fold, *q <* 0.05, independent of diurnal variation). Because effect-size and significance estimates differ systematically in scale and distribution between assay types, significant candidates were ranked by a scale-invariant *t*-statistic (the fatigue-response coefficient divided by its standard error) and capped at the 5 highest-ranked candidates per fold, rather than applying a single magnitude threshold across both assay types. As with the individual targeted and protein panels, the proportion of folds in which each candidate was retained in this joint screen was calculated, and the final combined biomarker panel was defined as those candidates retained in at least 80% of folds.

#### Temporal Analysis of Targeted and Protein Biomarker Concentrations

To characterize how biomarker concentrations changed across the full three-day protocol, linear mixed-effects models were fit for each of the six targeted biomarkers and for final retained proteins, with timepoint as a fixed effect and participant as a random intercept to account for repeated measures within individuals. Timepoint was treated as a categorical factor with Measure 3 (morning of day 2) as the reference level, as this represented the most proximal resting measurement to the fatigue protocol. Prior to model fitting, concentrations (or abundances for panel proteins) were log-transformed to address positive skew, which was confirmed via visual inspection of quantile-quantile plots of model residuals. Pairwise comparisons between timepoints were conducted with Tukey adjustment for multiple comparisons. To aid interpretability, significant pairwise contrasts were restricted to 5 physiologically meaningful comparisons: Baseline stability (Measure 1 vs. 3), diurnal effect (Measure 1 vs. 2), fatigue response (Measure 3 vs. 4), immediate recovery (Measure 4 vs. 5), and next day recovery (Measure 4 vs. 6 or 7).

#### Model Performance Comparison

As stated prior, ridge-penalized logistic regression was used to classify acute fatigue (Measure 3 vs. Measure 4) for all three biomarker panels (targeted, proteins, and combined), using the glmnet package (version 4.1-10) in R (version 4.5.2). Within each training fold, the ridge regularization strength (*λ*) was selected via an inner 5-fold cross-validation using the one-standard-error rule (`lambda.1se`) to favor a more parsimonious, less overfit model. Because logistic regression produces calibrated class probabilities directly via its link function, no additional probability calibration step was required. To directly compare the discriminative performance across these models, sensitivity, specificity, overall accuracy, F1 score, Matthews Correlation Coefficient (MCC), and area under the receiver operating characteristic curve (AUC) were calculated for each model across the nested LOSOCV folds, using each fold’s independently-selected panel.

Additionally, to evaluate the discriminative performance of the biomarkers most consistently identified across folds, the same classification metrics were calculated using each model’s final panel held fixed across all folds, rather than being allowed to vary by fold (fixed-panel classification performance). This allowed for quantification of fixed-panel classification performance which reflects how the panel would be expected to perform if applied consistently across new individuals rather than re-selected for each one, while also carrying a mild risk of residual information leakage relative to the fully nested classification performance above. Therefore, these results are reported as a secondary robustness analysis characterizing the discriminative value of the most consistently-identified candidates, rather than as an independent estimate of generalization performance. Permutation importance was calculated for the combined biomarker panel to assess each biomarker’s relative contribution to model performance. A ridge-penalized logistic regression was fit to the full sample and baseline classification accuracy was recorded. For each biomarker, values were randomly permuted across observations (1,000 iterations) and the fitted model was reapplied to the permuted data without refitting; permutation importance was defined as the mean reduction in classification accuracy across iterations relative to the unpermuted baseline, with larger reductions indicating greater reliance on that biomarker for classification.

To explore the temporal generalizability of each model beyond the training window, each folds’ trained ridge logistic regression model was applied to the corresponding heldout participant’s data under the nested framework, across all seven timepoints for the full three-day protocol. Standardization parameters were derived exclusively from the training participant’s Measures 3 and 4 values within each fold and applied consistently to all seven timepoints. At each time-point, participants were classified as Fatigued or Rested using the same 0.5 probability threshold as the primary classification analyses, and the percentage of held-out participants classified as Fatigued was calculated, with 95% Wilson score confidence intervals reflecting the binomial nature of this proportion.

## Results

### Change in Performance Parameters

Physical fatigue was defined as a categorical state of reduced physical performance, confirmed by significant declines in aerobic and anaerobic capacity, and muscular endurance following the protocol. Mean and standard deviation of performance measures pre- and post-fatigue protocol are presented in Table 1. Post-fatigue, average HR significantly increased by 69 bpm (*p* = 0.001) and recovery HR significantly increased by 32 bpm (*p <* 0.001), while average HRV significantly decreased by 65 ms (*p* = 0.008), confirming a significant cardiorespiratory response to the protocol. Muscular endurance measures were also significantly reduced, with maximum pushups declining by 6 repetitions (*p <* 0.001) and maximal plank hold duration decreasing by 57 s (*p <* 0.001). Anaerobic capacity also showed a significant decrease of 58 W following the protocol (*p* = 0.002). No significant changes were observed for maximum vertical leap (*p* = 0.617) or peak isometric force (*p* = 0.258).

**Table 1.** Changes in physical performance parameters pre- to post-fatigue protocol.

| Performance Parameter | Pre-Fatigue Mean (SD) | Post-Fatigue Mean (SD) | Effect Size (Hedges' *g*) | *p*-value |
| --- | --- | --- | --- | --- |
| **Cardiorespiratory Responses** |  |  |  |  |
| Heart Rate (bpm) | 67.2 (6.47) | 136.4 (21.4) | 2.57 | 0.001 |
| Heart Rate Variability (ms) | 86.5 (33.1) | 21.1 (10.8) | −1.44 | 0.008 |
| Recovery Heart Rate (bpm) | 128.7 (23.7) | 160.3 (18.46) | 1.84 | <0.001 |
| **Neuromuscular Performance** |  |  |  |  |
| Max Vertical Leap (cm) | 37.0 (9.6) | 37.5 (10.0) | 0.15 | 0.617 |
| IMTP Peak Vertical Force (N) | 2334.7 (1056.7) | 2223.2 (868.2) | −0.35 | 0.258 |
| Max Hand Release Pushups (#) | 38.2 (6.6) | 32.1 (6.6) | −1.40 | 0.001 |
| Max Plank (s) | 150.0 (48.9) | 92.9 (24.6) | −1.71 | <0.001 |
| **Metabolic Performance** |  |  |  |  |
| Anaerobic Capacity (W) | 448.0 (133.9) | 389.7 (125.3) | −1.24 | 0.002 |

*Note.* Values are mean (SD); effect sizes are Hedges' *g* for paired comparisons.

### Identified Targeted Biomarker Panel

Retained targeted markers in the nested classification procedure varied slightly across folds. IgA and UA were retained in all 10 folds, reflecting a consistent fatigue response and absence of diurnal variation across training subsets. AA was only retained in half of folds (5 of 10) with testosterone (2/10) and IL-6 (1/10) being retained in only a minority of folds, reflecting borderline effects to fatigue sensitive to which training subjects were included. Cortisol was not retained in any fold, consistent with the absence of any measurable fatigue response in the full-sample analysis. Applying the pre-specified 80%-of-folds threshold, the final targeted biomarker panel used for classification therefore consisted of only IgA and UA.

### Identified Protein Panel

Retained proteins in the nested classification procedure also varied across folds. Complement C1s subcomponent preproprotein (C1S) was retained in all 10 folds, reflecting a consistent fatigue response and absence of diurnal variation across training subsets. N-acetylmuramoyl-L-alanine amidase (XP_006722696.1; gene symbol not annotated), 26S proteasome regulatory subunit 6A (XP_016873515.1; gene symbol not annotated), and eosinophil peroxidase preproprotein (EPX) were retained in only 1/10 folds. Applying the pre-specified 80%-of-folds threshold, the final protein panel used for classification therefore consisted of only C1S.

### Imputation Sensitivity Analysis

Protein selection was robust to the choice of detection-limit substitution fraction. The number of proteins meeting the BH-corrected fatigue-significance criterion decreased monotonically as the substitution fraction increased (50%: 13 proteins; 75%: 10 proteins; 100%: 7 proteins), with strong pairwise overlap across all three (Jaccard = 0.54–0.77). Critically, each stricter substitution fraction identified a complete subset of the proteins identified at 50%, with no contradictory or reversed findings. Both proteins retained in the final nested cross-validation classification panels (C1S and ANG) were significant under all three substitution fractions tested, indicating that these specific biomarker findings are not sensitive to this modeling choice.

### Identified Combined Panel

Retained candidates in the nested classification procedure, in which targeted markers and proteins were screened jointly rather than separately, also varied across folds. IgA, UA, and C1S, each already identified as fully consistent (10/10 folds) in the separate targeted and protein panels above, were again retained in all 10 folds under joint screening, providing independent corroboration of their fatigue-response consistency. Angiogenin precursor (ANG), a protein not initially identified by the protein-only screen, was also retained in all 10 folds. Additionally, vesicle-associated membrane protein 7 (VAMP7) was retained in 2/10 folds, and receptor-type tyrosine-protein phosphatase C precursor (PTPRC), gelsolin isoform a precursor (GSN), protein ABHD14B (ABHD14B), keratin, type II cytoskeletal 6C (KRT6C), myeloperoxidase precursor (MPO), IL-6, prothrombin preproprotein (F2), and one unmapped protein (integrin alpha-M isoform 1 precursor) were each retained in only 1/10 folds. Applying the pre-specified 80%-of-folds threshold, the final combined panel used for classification therefore consisted of IgA, UA, C1S, and ANG.

### Temporal Responses of Targeted Biomarkers and Identified Proteins to Fatiguing Exercise

Mixed-effects models revealed distinct temporal patterns across biomarkers that varied in their relationship to fatigue status, diurnal rhythm, and recovery. No targeted biomarker or protein differed significantly between Measure 1 and Measure 2 or Measure 1 and Measure 3 (all *p >* 0.05), suggesting that pre-fatigue concentrations on day 2 were stable relative to the morning of day 1 and that limited diurnal effects were observed. With respect to the acute fatigue response, IgA (*p <* 0.001), UA (*p <* 0.001), AA (*p* = 0.020), IL-6 (*p* = 0.049), C1S (*p <* 0.001), and ANG (*p <* 0.001) were all significantly elevated at Measure 4 relative to Measure 3, whereas cortisol and testosterone did not show a significant fatigue response.

Recovery trajectories also differed across markers. IgA returned toward baseline within one hour post-protocol (*p <* 0.001), as did C1S (*p <* 0.001), ANG (*p* = 0.018), and testosterone (*p* = 0.017). By the following day (day 3), all markers that showed a significant fatigue response had returned toward pre-fatigue levels, with Measures 6 and 7 significantly lower than Measure 4 (all *p <* 0.05), with the exception of ANG which likewise declined significantly by Measure 6 (*p <* 0.001), but did not differ significantly from Measure 4 at Measure 7. Notably, cortisol showed significant suppression at Measure 7 relative to Measure 4 (*p <* 0.001), likely reflecting continued diurnal decline rather than a fatigue-specific recovery effect. Individual and group mean log-transformed concentrations (or abundances) of all retained candidate salivary biomarkers of acute fatigue (UA, IgA, C1S, and ANG) across all seven collection timepoints are shown in Figure 2.

**Fig. 2.**
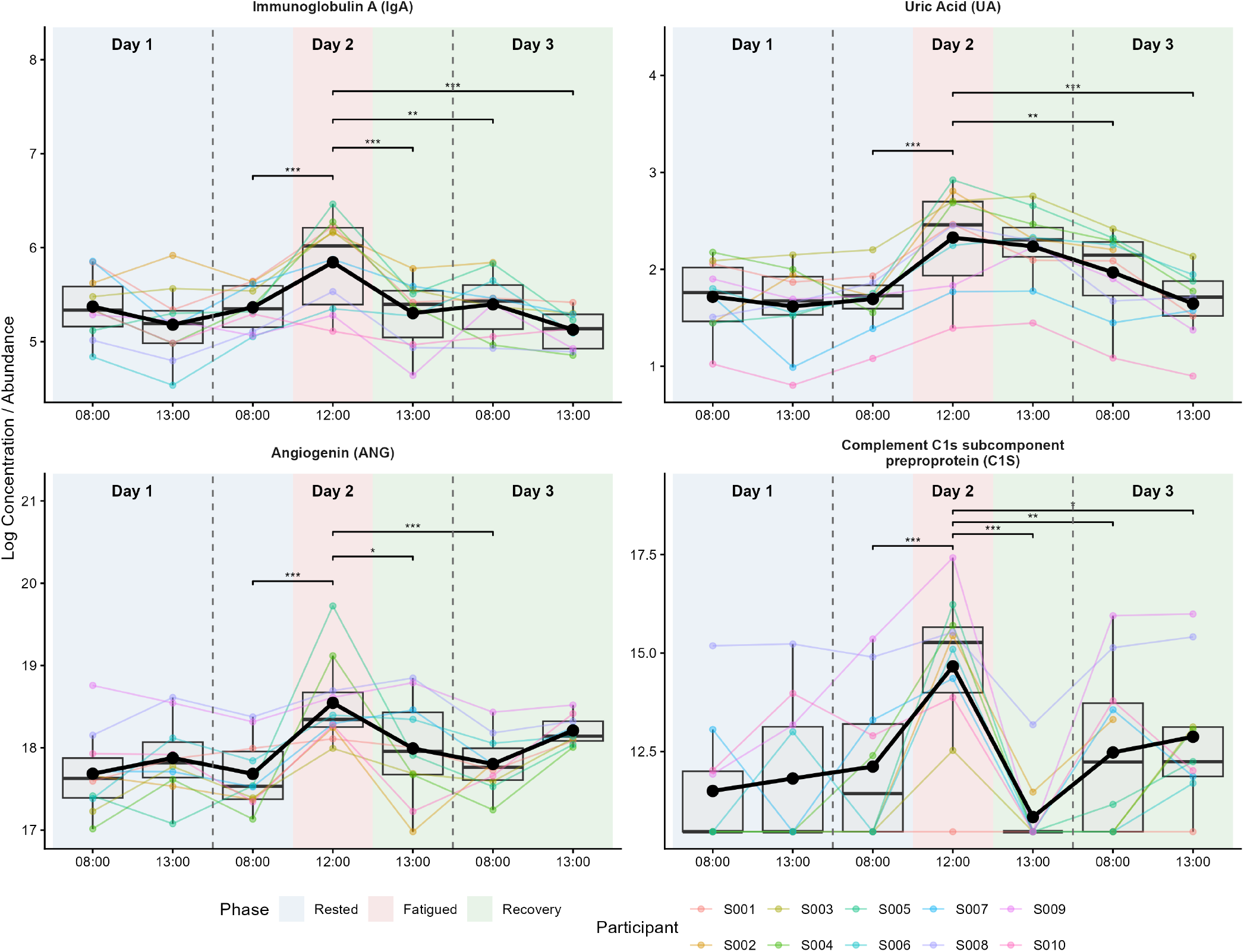
Log-transformed concentrations/abundances of identified panel salivary biomarkers across the three-day protocol. The bold line represents group means at each timepoint and colored lines represent individual participant trajectories. Shaded regions denote protocol phases: Rested (Measures 1–3; blue), Fatigued (Measure 4; red), and Recovery (Measures 5–7; green). Dashed vertical lines and day labels demarcate the three-day collection schedule while x-axis labels indicate the approximate clock time of each saliva collection. Significance brackets indicate pairwise differences surviving Tukey correction (*p <* 0.05), restricted to five physiologically meaningful comparisons: Baseline Stability (Measure 1 vs. 3), Diurnal Effect (Measure 1 vs. 2), Fatigue Response (Measure 3 vs. 4), Immediate Recovery (Measure 4 vs. 5), and Next Day Recovery (Measure 4 vs. 6 or 7). \**p <* 0.05, \*\**p <* 0.01, \*\*\**p <* 0.001.

### Comparison of Model Performance

Table 2 presents model performance metrics for both nested LOSOCV and fixed-panel LOSOCV frameworks across panels (targeted, proteomic, and combined). Under the nested framework, the targeted biomarker model correctly classified 6/10 fatigued and 8/10 rested participants, for an overall accuracy of 70% and AUC of 0.76. The protein model (C1S only) performed comparably to, or modestly better than, the targeted biomarker model under this framework, correctly classifying 8/10 fatigued and 7/10 rested participants, for an overall accuracy of 75% and AUC of 0.76, reflecting higher sensitivity but somewhat lower specificity than the targeted model. The combined model, including IgA, UA, C1S, and ANG outperformed both single-platform nested models, correctly classifying 9/10 fatigued and 9/10 rested participants, for an overall accuracy of 90% and an AUC of 0.90.

**Table 2.**
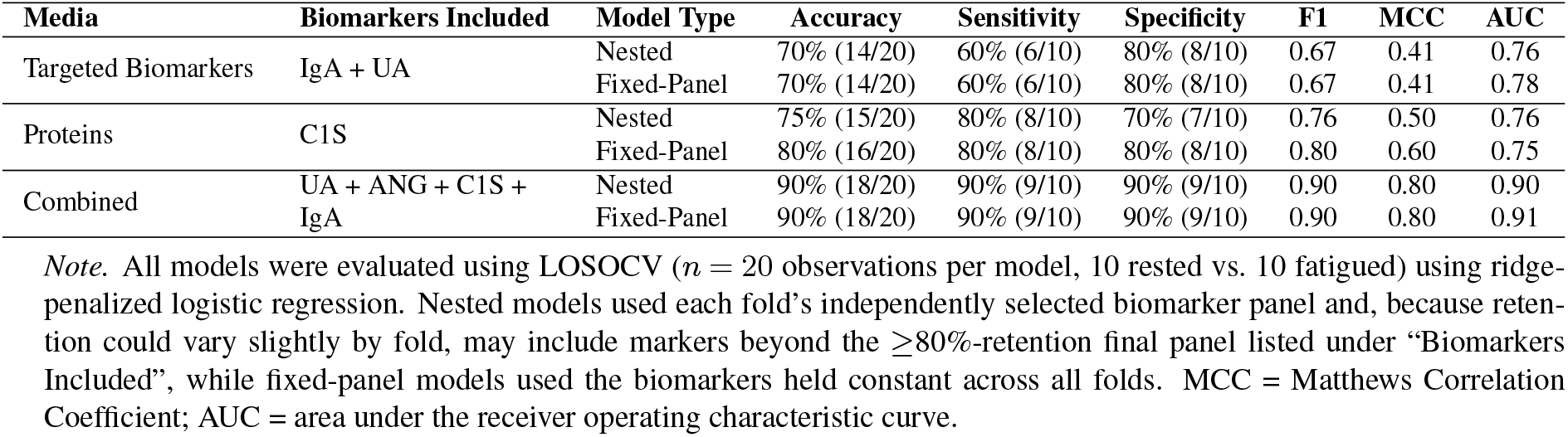
Comparison of model classification performance across media (targeted biomarkers, proteins, and combined) and model type (nested LOSOCV vs. fixed-panel LOSOCV).

Fixed-panel performance was closely aligned with the nested estimates across all three panels. For the targeted biomarker panel, fixed-panel performance was virtually identical to the nested estimate (60% sensitivity, 80% specificity, 70% accuracy), differing only marginally in AUC (0.78 vs. 0.76). For the protein-only panel, restricting classification to the single consistently-retained protein (C1S) modestly improved performance relative to the nested estimate and outperformed the fixed-panel targeted biomarker model, correctly classifying 8/10 fatigued and 8/10 rested participants, with a corresponding increase in accuracy (75% to 80%), MCC (0.50 to 0.60), and F1 score (0.76 to 0.80). The combined panel achieved the strongest classification performance among all model-panel combinations under both designs, correctly classifying 9/10 fatigued and 9/10 rested participants in each case (overall accuracy of 90%), with identical F1 (0.90) and MCC (0.80) and only a marginally higher AUC under the fixed-panel design (0.91 vs. 0.90). This close correspondence between the nested and fixed-panel estimates suggests the identified biomarker signatures reflect stable, reproducible fatigue signals rather than estimates inflated by the mild residual leakage inherent to the fixed-panel design. Permutation importance indicated that no single biomarker dominated the combined model’s fatigue classification. IgA contributed the most to model accuracy (permutation importance = 0.139), followed by ANG (0.116), C1S (0.103), and UA (0.096), suggesting the model’s performance reflects complementary information from both targeted and proteomic assay types rather than reliance on a single biomarker.

Predicted fatigue classification rates across the full 3-day protocol are shown in Figure 3. All three models classified comparatively few participants as fatigued during the rested phase (Measures 1–3), with the combined panel showing the lowest and most consistent rate (0–10% of participants classified as fatigued) relative to the targeted biomarker (10–20%) and protein-only (20–30%) models. At Measure 4 (immediately post-fatigue protocol), the combined panel classified the highest percentage of participants as fatigued (90%, 9/10), followed by the protein-only panel (80%, 8/10) and the targeted biomarker model (60%, 6/10). During the recovery phase, all three models showed a decline in the percentage of participants classified as fatigued from Measure 4 to 5 with the combined (90% to 20%) and protein-only panels (80% to 20%) showing a much greater reduction relative to the targeted panel (60% to 50%). The protein-only model rose again relative to Measure 5 at Measure 6 (40%) before declining at Measure 7 (22%), while the combined panel remained low at Measure 6 (10%) before a modest rise at Measure 7 (33%) while the targeted biomarker model declined more gradually across Measures 6 and 7 (40% and 11%, respectively). Ninety-five percent Wilson score confidence intervals were wide throughout, reflecting the limited precision of proportion estimates at this sample size (*n* = 9–10 participants per timepoint), and should be considered when interpreting between-panel and between-timepoint differences.

**Fig. 3.**
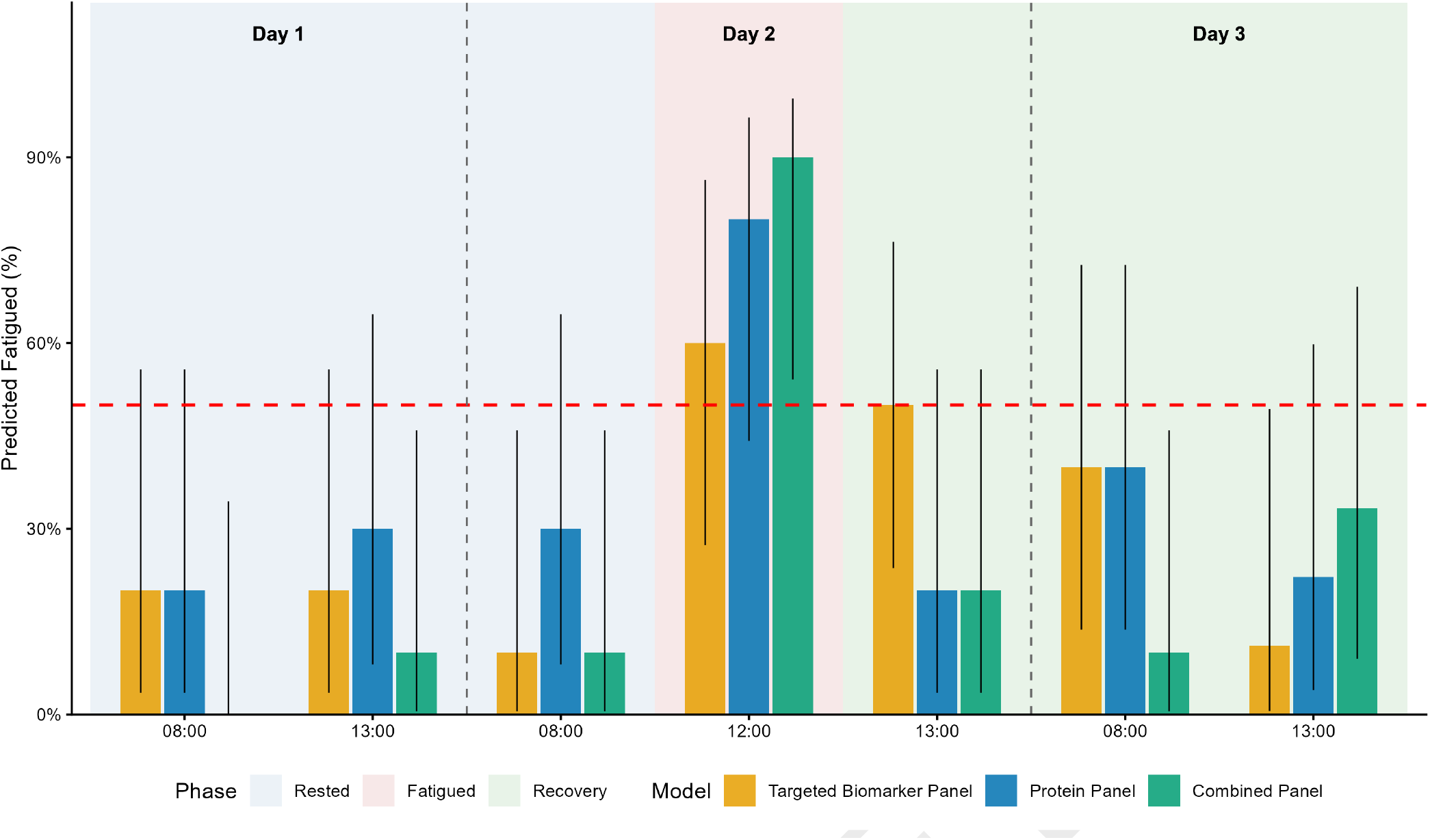
Predicted fatigue classification rate trajectories for the targeted biomarker, salivary protein, and combined models applied across all seven collection timepoints of the three-day protocol. Bars represent the percentage of participants classified as fatigued (predicted probability *≥* 0.50) at each timepoint, with error bars indicating the 95% Wilson score confidence interval. The red dashed line denotes the 0.50 classification decision boundary. Background shading indicates protocol phase: Rested (Measures 1–3; blue), Fatigued (Measure 4; red), and Recovery (Measures 5–7; green).

## Discussion

Salivary biomarkers offer potential as a non-invasive way to monitor physiological stress, with prior studies linking salivary biomarkers to fatigue and performance outcomes in tactical and athletic settings (23–25). This study builds on this work by identifying and comparing panels of salivary biomarkers that are capable of classifying acute physical fatigue in an exploratory sample: one based on targeted stress-related biomarkers, another on a candidate protein identified through data-driven proteomic profiling, and a third integrating markers from both approaches into a combined, multi-platform panel. Using the fixed biomarker panels identified through nested cross-validation, both single-platform models achieved moderate classification accuracy with the protein-only model correctly classifying more fatigued observations than the targeted biomarker model (8/10 vs. 6/10) and slightly more observations overall (16/20 vs. 14/20) while the two models classified an equal number of rested observations correctly (8/10 each). The combined model, consisting of two targeted salivary biomarkers (IgA and UA) and two salivary proteins (C1S and ANG), outperformed both single-platform models, correctly classifying 18/20 observations overall, including 9/10 fatigued and 9/10 rested observations.

Cortisol and dehydroepiandrosterone (a precursor to testosterone) have been studied as potential biomarkers for stress resilience during military training (26, 27); however, in the present study neither cortisol nor testosterone showed a significant fatigue-specific response (Measure 3 vs. Measure 4), either in the full sample or consistently across LOSOCV folds. Cortisol was not retained in any of the 10 nested folds, while testosterone was retained in only 2/10, indicating that any apparent fatigue association for testosterone was unstable rather than robust. These findings are consistent with the broader literature on these hormones as both cortisol and testosterone are known to respond broadly to psychological, social, and circadian influences well beyond physical fatigue itself, and both exhibit substantial diurnal variation that can obscure a fatigue-specific signal (26, 27). Individual variability in cortisol reactivity may be further compounded by genetic differences in glucocorticoid receptor sensitivity (28) and by the disproportionate influence of psychological, relative to physical, stressors on cortisol secretion (16). As our screening procedure explicitly required a fatigue-specific response independent of diurnal (morning-to-afternoon) variation, the well-documented lack of specificity in cortisol and testosterone likely explains their exclusion from the final panel in favor of biomarkers with more fatigue-specific response profiles.

AA and IL-6, both of which are linked to sympathetic activity and inflammation (9, 10), showed smaller but significant fatigue responses (AA: *p* = 0.020; IL-6: *p* = 0.049); however, both failed to be robustly selected across LOSOCV folds (AA: 5/10, IL-6: 1/10), indicating that these full-sample effects, while statistically significant, were not sufficiently stable or precise to survive resampling at this sample size. This instability may be attributable to variability in secretion during fatigue, which may be influenced by differences in hypothalamic-pituitary-adrenal (HPA) axis responsiveness (29), genetic predispositions (30), anticipatory stress responses (31), hydration status (32), and baseline psychological or physiological stress levels (33), potentially limiting their discriminative sensitivity for acute fatigue detection.

Within the retained targeted biomarker panel, UA and IgA demonstrated the most statistically robust response to acute fatigue, surviving selection in 100% of nested LOSOCV folds. The acute elevation of UA is consistent with its established role as a marker of purine nucleotide degradation and metabolic turnover during high-intensity exertion as intense, glycolytically demanding exercise depletes muscle ATP and accelerates purine catabolism to uric acid (34). Uric acid’s antioxidant properties may further contribute to its reliability as a fatigue marker, as circulating UA is oxidized in skeletal muscle during high-intensity exercise and has been shown to blunt exercise-induced oxidative stress in healthy adults (35). IgA’s response is consistent with its role in mucosal immune defense, which increases with moderate exercise in sedentary adults (8), a pattern broadly consistent with the elevation observed here. Our findings, together with those in prior literature, suggest that the markers retained in the final targeted panel may reflect more stable, mechanistically direct responses to acute physical fatigue than the more broadly stress-reactive or context-dependent hormonal and enzymatic markers evaluated above.

Only one protein, C1S, was robustly selected into the final protein-only panel. Despite representing a single-marker panel, C1S demonstrated better overall accuracy and sensitivity than the multi-marker targeted panel under the same conservative cross-validation framework, though the two panels showed comparable AUC. The robustness with which C1S was selected across such a small, exploratory sample, paired with its classification performance, provides evidence for it as a candidate salivary biomarker of acute physical fatigue worthy of further investigation in larger samples. C1S, the serine protease subunit of the C1 complement complex, initiates the classical complement pathway and reflects rapid innate immune activation in response to tissue stress and cellular damage (36). Its elevation in saliva following exertion is consistent with prior evidence that strenuous exercise activates classical complement pathway markers, including C1s, particularly during resistance exercise and field-based protocols designed to elicit muscle damage (37, 38). This pattern extends to other field-based salivary proteomic investigations of military and occupational stress where immune-related proteins have similarly emerged as a dominant molecular signature (14, 15). Complement and coagulation proteins, including multiple subcomponents of the C1 initiator complex, were among the most strongly upregulated clusters during chronic military stress (15), and immunoregulation proteins comprised the largest functional category among fatigue-related salivary proteins identified in emergency physicians (14). These findings are consistent with the identification of C1S as the most robustly selected protein marker of acute fatigue in the present study, though direct comparison is limited by differences in fatigue-induction protocol (sustained military/occupational stress vs. acute physical exertion) and proteomic methodology between studies.

The identification of a combined, multi-platform panel, comprised of two targeted stress markers (IgA, UA) and two salivary proteins (C1S, ANG), is consistent with a broader shift toward multi-domain biomarker approaches in military and occupational physiology research. Integrating multiple physiological inputs may allow for a more complete characterization of complex outcomes such as acute fatigue, which likely reflects the combined influence of immune, metabolic, and cellular stress processes rather than any single pathway (4). This approach may also provide greater robustness to potential confounders, such as diurnal rhythm, that can limit the reliability of single-analyte predictions (27), and reflects a growing recognition that integrating multiple molecular data streams provides a more comprehensive understanding of the biological response to physiological stress than any single assay type alone (15).

While IgA, UA, and C1S have each been discussed above, ANG’s inclusion in the combined panel is, to our knowledge, a novel finding in the context of acute physical fatigue. Angiogenin (ANG) is a multifunctional ribonuclease with an established role in the cellular stress response, translocating to the cytoplasm under oxidative, nutrient, and mitochondrial stress to generate stress-induced tRNA fragments that support cell survival (39). Although this mechanism has not previously been examined in the context of acute physical fatigue, it provides a plausible biological basis for ANG’s elevation at peak fatigue observed here, reflecting cellular-level stress signaling that is mechanistically distinct from, and complementary to, the innate immune activation reflected by C1S. Notably, ANG was not identified by the protein-only screen despite being retained in all 10 folds of the joint screen reflecting the methodological difference between the two procedures with the joint screen ranking candidates by a scale-invariant *t*-statistic and retaining only the top five per fold, whereas the protein-only screen applies an absolute effect-size threshold. This ranking-based approach can surface a candidate with a precise but comparatively modest effect size that would not clear the fixed threshold used when proteins are screened in isolation, therefore, ANG’s absence from the protein-only panel may reflect the stringency of that threshold rather than evidence against a genuine, if comparatively modest, fatigue-related signal. Considered together, the four biomarkers in the combined panel span distinct but interconnected levels of the physiological stress response with IgA reflecting sympathetically-mediated mucosal immune activation, UA reflecting metabolic turnover and antioxidant defense against oxidative stress, C1S reflecting tissue-level innate immune activation in response to cellular damage, and ANG reflecting intracellular stress signaling.

Beyond classification accuracy at a single fatigue timepoint, we examined each panel’s predicted-fatigued classification rate across the full seven-timepoint collection window to evaluate how well each panel captured the trajectory of rested state to fatigue onset and finally recovery. The divergence in fatigue classification rate during recovery (Measures 5– 7) between the targeted, protein, and combined panels may reflect a combination of differences in marker-specific recovery kinetics and a limitation of the panel-selection procedure itself as feature selection was based solely on a fatigue-specific contrast (Measure 3 vs. Measure 4) and a diurnal-specificity criterion (Measure 1 vs. Measure 2). Therefore, no stage of screening selected for stability during recovery, leaving each panel’s post-fatigue behavior effectively unconstrained by the selection process. The targeted panel showed the clearest, most gradual decline in fatigue classification percentage from Measure 4 to 7, a pattern with some biological grounding for UA specifically, as it has been shown to accumulate with a delayed time course following intense exercise and remains elevated as an antioxidant response (40). Salivary IgA’s recovery kinetics are considerably less consistent in the broader literature (41), which may have contributed to the targeted panel’s own residual variability. Both the protein-only and combined panels’ advantage in correctly recognizing the rested state at Measures 1–3 was not matched by comparable stability during recovery, with fatigue classification sharply dropping at Measure 5 for both followed by a rebound in the protein-only panel at Measure 6 and for the combined panel at Measure 7, suggesting that the overall reliability is concentrated in fatigue onset and rest for the protein-containing panels, rather than sustained through the full physiological timeline. However, without concurrent objective performance testing or subjective readiness assessments collected at each recovery timepoint in a larger sample, it remains unclear which panel’s post-fatigue trajectory most accurately reflects true physiological recovery, as opposed to reflecting noise inherent to a small-sample, threshold-based classification metric.

Pre-post differences in physiological measures and performance assessments confirmed that the fatiguing protocol elicited a high level of acute physical fatigue in our sample. Following the protocol, heart rate increased by an average of 69 bpm (*∼* 103%), while HRV declined by 65 ms (76%), reflecting a marked shift toward sympathetic dominance consistent with substantial physiological stress. These autonomic responses were accompanied by meaningful reductions in muscular endurance, anaerobic capacity, and aerobic recovery performance, collectively confirming that the protocol successfully induced acute fatigue. In contrast, maximum vertical jump height did not significantly change, suggesting that the protocol produced substantial physiological and performance strain without impairing gross explosive lower-body performance as assessed by jump height alone. This finding is consistent with load carriage literature showing that lower-body power outcomes do not always decline following military or occupationally relevant tasks (42). A systematic review by Sax van der Weyden et al. (2026) reported that some studies have found reduced countermovement jump height after foot march exercise, while several others have found no significant change in countermovement jump height or broad jump performance following loaded foot marches (42). One possible explanation is that the protocol induced greater autonomic and metabolic disturbance than neuromuscular impairment detectable by jump height alone. Additionally, because participants completed the protocol while wearing a weighted vest, removal of that load prior to post-testing may have contributed to a transient post-activation potentiation effect that offset fatigue-related decrements (43). The overall magnitude and pattern of these responses are broadly consistent with those reported following military-relevant physical tasks, including decrements in aerobic and anaerobic capacity and muscular performance following short-duration field exercises (44), suggesting that the protocol reasonably approximated the physiological demands of occupational physical activity, albeit in a recreationally active rather than tactical sample.

### Strengths, Limitations, and Implications for Future Research

Although derived from a small, exploratory sample, our results suggest that salivary proteomic profiling has potential to objectively classify acute physical fatigue status relative to traditionally studied stress biomarkers, and that multi-platform profiling may offer additional classification benefit beyond either single-platform approach alone. While this untargeted, data-driven approach offers a promising starting point for identifying candidate proteins associated with acute physical fatigue, the large number of quantified proteins relative to the limited sample size introduces a high risk of overfitting and inflated effect estimates. To directly address this, protein screening incorporated a Stage-0 reliability pre-filter (restricting candidates to the most measurement-consistent half of proteins by rested-phase variability), within-fold Benjamini-Hochberg false discovery rate correction, and an 80%-of-folds stability-selection threshold for final panel inclusion. Alongside a conservative nested LOSOCV approach used on a balanced sample of immediately pre- and post-fatigue observations to best identify robust candidate markers, we attempted to balance the constraints of a small sample with the temporal resolution of our data collection design. Despite this, these results must be treated as exploratory and considered as hypothesis-generating pending validation in larger, independent cohorts before any clinical or operational application.

Additionally, the short protocol focused on acute fatigue and did not capture the chronic fatigue commonly experienced during extended training or deployment. Chronic fatigue arises from different mechanisms such as persistent inflammation (45), mitochondrial dysfunction (46), and hormonal dysregulation (47) which were not evaluated. Future studies should incorporate multi-day or cumulative workload designs to better reflect operational demands (44). Lastly, expanding to multi-omics approaches, including metabolomics, genomics, and transcriptomics, could provide a more integrated view of fatigue biology. Combining these with wearable monitoring, and considering sex-specific responses and individual variability, would improve model precision. Ultimately, translating these findings into targeted interventions could support effective fatigue management in high-demand environments.

## Conclusions

This exploratory study demonstrated that a combined, multi-platform panel integrating targeted salivary biomarkers and candidate salivary proteins of acute fatigue outperformed either single-platform approach in classifying acute physical fatigue in a small, recreationally active sample, while a single candidate protein, C1S, identified through untargeted proteomic screening, achieved classification performance comparable to a multi-marker targeted panel on its own. These preliminary findings support greater examination of proteomic and multi-platform approaches for quantifying physical readiness within tactical and occupational athlete populations where traditional performance and/or readiness assessments may not be feasible.

### Practical Applications

Findings from this exploratory study suggest that salivary biomarkers may offer a practical method for monitoring acute physical fatigue that can be extended to occupational and tactical settings. In particular, C1S emerged as a candidate salivary biomarker of acute physical fatigue, while the combined, multi-platform panel showed the most promise for accurate physical fatigue profiling, suggesting that a combinatorial approach could best inform training load management, fatigue risk mitigation, and operational readiness decisions.

## Acknowledgments

N/A

## Data Availability

The data and analysis code that support the findings of this study are openly available. The complete analysis pipeline, targeted biomarker and proteomic datasets, and all reported results are publicly available at https://github.com/blindse3/salivary-biomarkers-fatigue.

## Disclaimer

Material has been reviewed by Walter Reed Army Institute of Research. There is no objection to its presentation and/or publication. The opinions or assertions contained herein are the private views of the author, and are not to be construed as official, or as reflecting the views of the Department of the Army or the Department of Defense.

